# Differential coping strategies exerted by biofilm and planktonic cells of the beneficial bacterium *B. subtilis* in response to the protozoan predator *Entamoeba histolytica*

**DOI:** 10.1101/2024.11.16.623985

**Authors:** Ilana Kolodkin-Gal, Prem Anand Murugan, Eva Zanditenas, Serge Ankri

## Abstract

The human protozoan parasite *Entamoeba histolytica* causes amebiasis and interacts with both beneficial and harmful members of the microbiome. In previous studies, it was shown that *E. histolytica* can break down pre-established biofilms of *B. subtilis* in a time- and dose-dependent manner. Inhibiting parasitic cysteine proteases impairs biofilm degradation. However, it is still unknown whether bacteria can sense this process and respond to the degradation of the biofilms.

Here, our research demonstrates a multi-layered response of probiotic bacteria to the parasite, which differs between planktonic bacteria and pre-established biofilms. Sensing the activity of cysteine proteases from *E. histolytica*, the bacteria activate the general stress response and, to a lesser extent, the cell wall stress response, making the surviving biofilm members more resistant to mild stressors. On the other hand, planktonic cells exposed to the predators’ lysate deactivate the expression of genes associated with biofilm formation while inducing their motility to avoid predation. Overall, our results indicate that bacteria have evolved to recognize amoeba predators. Furthermore, the partially digested biofilm cells may have unexpected disadvantages over bacteria that did not encounter a predator. These findings may be useful in developing more efficient probiotic strains that are resilient to amoebiasis.

## Introduction

*Entamoeba histolytica* is a protozoan parasite responsible for amebiasis, a wildly prevalent human intestinal disease in developing countries. Transmission of amebiasis occurs mainly through contaminated food or water with feces containing *E. histolytica* cysts, which is one of two forms of the parasite ^1^. On entering the host intestine, the cyst, the resistant form of the parasite, undergoes a process called excystation during which trophozoites, the vegetative form of the parasite is released^2^. In most of the cases of infection, these trophozoites feed on bacterial microbiota or cellular debris in the large intestine without causing symptoms.

*Bacillus subtilis* is a bacterium capable of forming biofilms. It is commonly found in soil and is used as a probiotic to support digestive and immune health in both adults and children. When grown in isolation, it forms robust biofilms both on liquid surfaces and agar plates, which are triggered by factors like low oxygen and nutrient depletion. The process is governed by the master regulator Spo0A. The initial step involves the transition of planktonic cells to a sessile state, with a decrease in flagella gene expression and an increase in genes responsible for producing the extracellular matrix (ECM)^3,4^. This matrix consists of extracellular polysaccharides and proteins, with TasA and TapA contributing to the 3D architecture of the biofilms ^5-9^ and are coregulated with an additional protein barrier, the hydrophobin BslA ^10,11^. TasA regulates gene expression in the complex biofilm structures ^12-14^ and promotes the formation of crystalline calcium carbonate ^11,15^.

In addition to biofilm formation, the transcriptional stress response in *B. subtilis* can be activated by the general stress response regulators, primarily the alternative sigma factor SigB ^16^. The activation of this sigma factor, transiently induced after the imposition of stress, allows *B. subtilis* to exhibit an immediate response to various stresses through the induction of over 150 general stress proteins (GSP)^16^.

Our previous research explored the molecular mechanisms involved in the interaction between parasites and biofilms that coexist in the same habitat, specifically the gut ^17^. We used genetically manipulable models of *E. histolytica* and *B. subtilis* for our study. Once bacteria form biofilms, simple phagocytosis becomes unfeasible, as the biofilms grow too large to be engulfed. Therefore, the degradation of the biofilms and release of smaller particles is essential for the amoeba to feed ^18^. We identified the unique transcriptome of *E. histolytica* after exposure to *B. subtilis* biofilm, the role of cysteine proteases in the degradation of the biofilm, and the protective role of the *B. subtilis* biofilm against oxidative stress. These results highlighted the differential response of the parasites to planktonic and biofilm cells ^17^. However, the response of the bacteria to the parasite, and whether it is different between planktonic cells and biofilm cells remained to be determined.

In this work, we deepen our understanding of the response of biofilm cells to the predator and the predation process as judged by their response to active CPs from the extract. Following predation, biofilm cells lose their tolerance to stressors due to the dissolution of the extracellular matrix. Our results indicate that biofilm cells sense CPs and activate the general stress response associated genes and a beta lactamase associated with the cell wall stress response, which allow them to survive moderate stressors and antibiotic concentrations upon release to the environment.

During liver infection, a massive death of trophozoites is observed in the first hours postinfection due to the activation of the immune response, releasing lysate content into the blood stream and subsequently to the GI^19^. Death of free living trophozoites may also occur during cell division and with exposure to disinfectant ^20^.

Hence, exposure to the parasite lysate may act as a warning signal for bacteria, prompting them to avoid forming biofilms and instead promoting movement. This ability enables them to effectively evade predation by moving away from potential predators. Interestingly, the stress response is activated in established biofilms, while the downregulation of matrix genes is more common in planktonic bacteria. Overall, our findings suggest that biofilm-forming cells have co-evolved with potential predators, enabling the prey to survive predation and coexist in the same environment as their predators.

## Methods

### Culture of *E. histolytica*

*E. histolytica* trophozoites, the HM-1: IMSS strain (from Prof. Samudrala Gourinath, Jawaharlal Nehru University, New Delhi, India), were grown and harvested according to a previously reported protocol ^21^.

### Extract Preparation

*E. histolytica* trophozoites (1 × 10^6^) total lysates were prepared using a lysis buffer containing 1% Nonidet P-40 (NP-40 in Deuterium-depleted water (DDW)). The trophozoites were incubated 15 minutes with NP-40 on ice and centrifuged for 10 minutes at 12 000 rpm at 4 degrees Celsius. The supernatant was then transferred to a new tube. Protein concentration was measured by Bradford.

### Strains and Media

All strains were derived from *B. subtilis* NCIB 3610^22^. A complete strain list is shown in Supporting Information Table S1.

The strains were routinely grown in LB broth (Difco) or MSgg medium^22^ (5 mM potassium phosphate, 100 mM MOPS (pH 7), 2 mM MgCl2, 50 μM MnCl2, 50 μM (liquid assay) and 125 μM (biofilm assay) FeCl3, 700 μM CaCl2, 1 μM ZnCl2, 2 μM thiamine, 0.5% glycerol, 0.5% glutamate, 50 μg/ml threonine, tryptophan, and phenylalanine). Solid medium contained 1.5% bacto agar (Difco).

### Flow Cytometry

Starter cultures were spotted on MSgg either applied or non-applied with amoeba extract. The plates were then incubated as mentioned in the first section. The plates were then incubated at 30 °C and biofilms colonies were harvested at 48 h post-inoculation and treated as discussed in each figure legends.

This was followed by sonication using a BRANSON digital sonicator. After sonication, samples were diluted in PBS and measured using an LSR-II cytometer (Becton Dickinson, San Jose, CA, USA). The GFP fluorescence was measured using laser excitation of 488 nm, coupled with 505 LP and 525/50 sequential filters. To distinguish background fluorescence from the reporters’ specific fluorescence, the WT *B. subtilis* grown under the same conditions was used as a negative control, and its background fluorescence was gated to separate the true fluorescent population (population outside the background gate) from the reporters as done in Maan et al. ^23^. A total of 100,000 cells were counted for each sample and flow cytometry analyses was performed using FACS Diva (BD biosciences) and FCS express 7 Research Edition.

### Statistics

All experiments were analyzed by GraphPad software (license to the Scojen Institute) with Anova test for multiple comparisons, unless stated otherwise.

## Results

### *B. subtilis* biofilm cells induce the general stress response following exposure to CPs

We previously demonstrated that *B. subtilis* biofilm cells, serve as prey for the *E. histolytica* trophozoites ^17^. This gradual degradation of the established biofilms could trigger alterations in gene expression ^4^. Specifically, it was shown that the protein TasA, degraded by the Parasite, is a regulator of the general stress response ^12^. Therefore, we evaluated the general stress response of the bacterium following exposure to amoeba extract. In short, we exposed pre-established biofilm cells to the extract for 30 minutes in increased doses of the extract and then evaluated their transcriptional response. To specifically monitor the general stress response, we assessed the transcription from the *ctc* Promoter, directly regulated by SigB ^24^ in biofilm cells before and following exposure to the amoeba signal. Our results demonstrated a significant dose-dependent induction of the *ctc* promoter (Fig. 1A).

**Figure 1:**
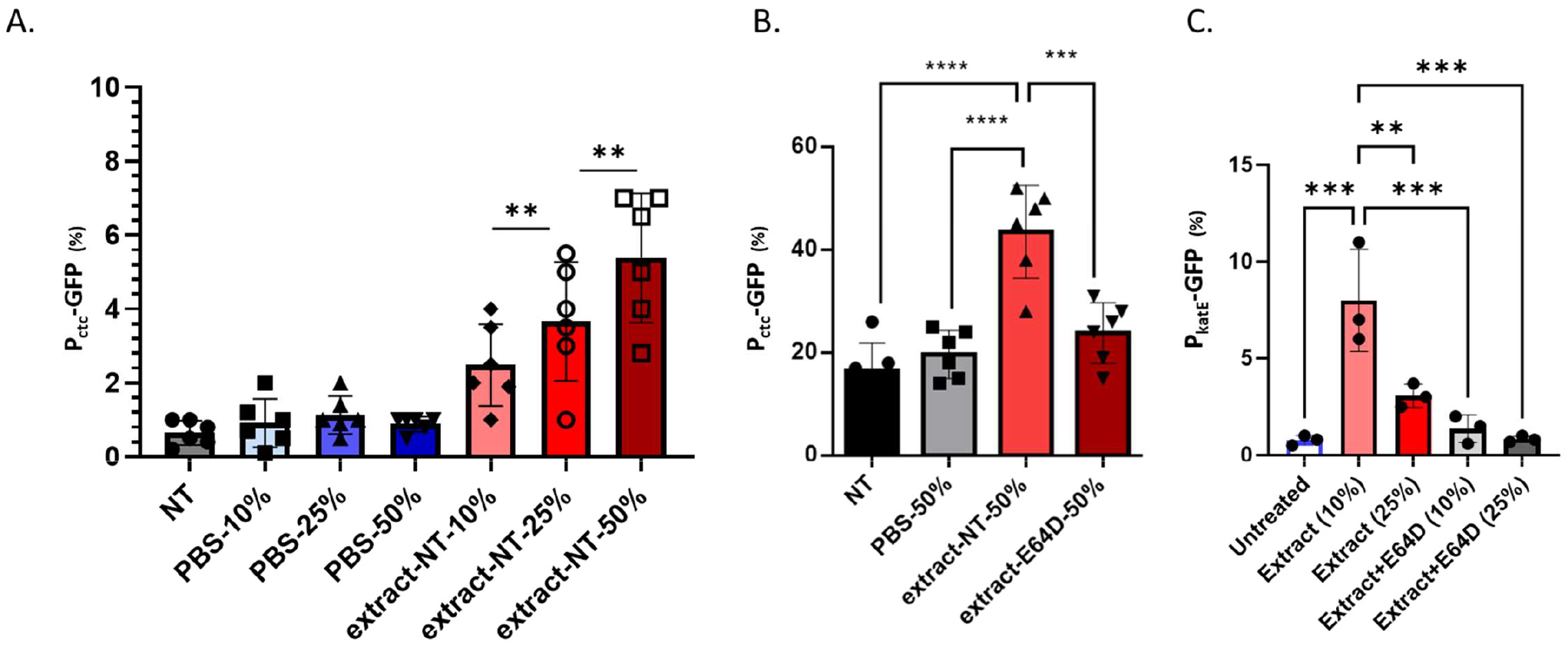
Amoeba CPs activity triggers the bacterial stress response. (A, B) *B. subtilis* strain NCIB3610 harboring *amyE::P*_*ctc-*_*GFP* (General stress response) was analyzed in the presence and absence of increasing concentrations of E. histolytica’s extract using flow cytometry. Colonies were grown on MSgg medium and incubated at 30 °C. To determine the effect of amoeba’s lyzate, the 48-hour biofilms were harvested. Colonies were incubated in MSgg containing the indicated concentrations of PBS/Indicated concentrations of extract/ Indicated concentrations of extract+E64D for 4 hours. Extracts were diluted v/v from a stock of 2ng/ μL. From untreated and treated biofilms, 100,000 cells were counted with flow cytometry. The % of cells expressing the reporters was calculated. Graphs represent mean ± SD from six independent experiments (n = 3). **<0.01, ***<0.001 and ****<0.0001. For B, extracts with and without E64D were used at identical concentrations (diluted v/v from a stock of 2ng/ μL). (C) *B. subtilis* NCIB3610 strain harboring *amyE::P*_*katE-*_*GFP* (General stress response) was analyzed in the presence and absence of increasing concentrations of *E. histolytica’s* extract using flow cytometry as indicated above.

We then asked whether this induction depends on the degradation of biofilms by amoeba CPs (Fig. 1A). We utilized the cysteine proteinases inhibitor E64D, which allows for the expression of the CPs by the parasite but prevents their activity and consequently biofilm and *tasA d*egradation ^17^. Our results indicated that extract with the chemical inhibitor for CPs activity E64D failed to induce the activation of SigB-dependent transcription (Fig 1B). This result suggests that the stress response is a readout of the biofilm degradation activity of the CPs rather than their mere presence.

The extract and extraction buffer were not toxic to pre-established biofilm cells at the tested concentrations (Figure S1). The buffer and E64D showed only mild inhibition of ctc transcription (Figure S2) and could not explain the observed stress induction. Thus, the activation of the stress response is specific and serves as a physiological indicator of biofilm degradation.

.To further confirm the specific response of SigB dependent promoters to *B. subtilis* biofilm degradation by amoeba CPs, we tested the transcription from an additional SigB dependent promoter, *katE* promoter ^25^ (encoding the a Catalase). This promoter is almost exclusively under SigB control in non-sporulation conditions. As shown, the exposure of biofilm cells to amoeba’s extract induced the transcription from *katE* promoter and was fully dependent on the presence of CPs (Fig. 1C). These results confirm that exposure to amoeba extract and subsequent biofilm degradation by CPs activate the general stress response in biofilm cells.

It was previously demonstrated that the *SigB* mutant impairs growth under ethanol and hydroxyl radical stress but does not affect growth under normal conditions. Therefore, we suspected that the induced stress response following an increased tolerance to hydroxyl radicals. Consistently, we found that the pre-established biofilm cells treated with amoeba extract were indeed more resistant to H_2_O_2_ (Fig. 2A) and this was dependent on the presence of Amoeba’s extract, and active CPs. Similar results were obtained with EtOH (Fig. 2A and 2B). Therefore, the activation of the general stress response during biofilm degradation is associated with elevated stress tolerance for mild general stressors.

**Figure 2:**
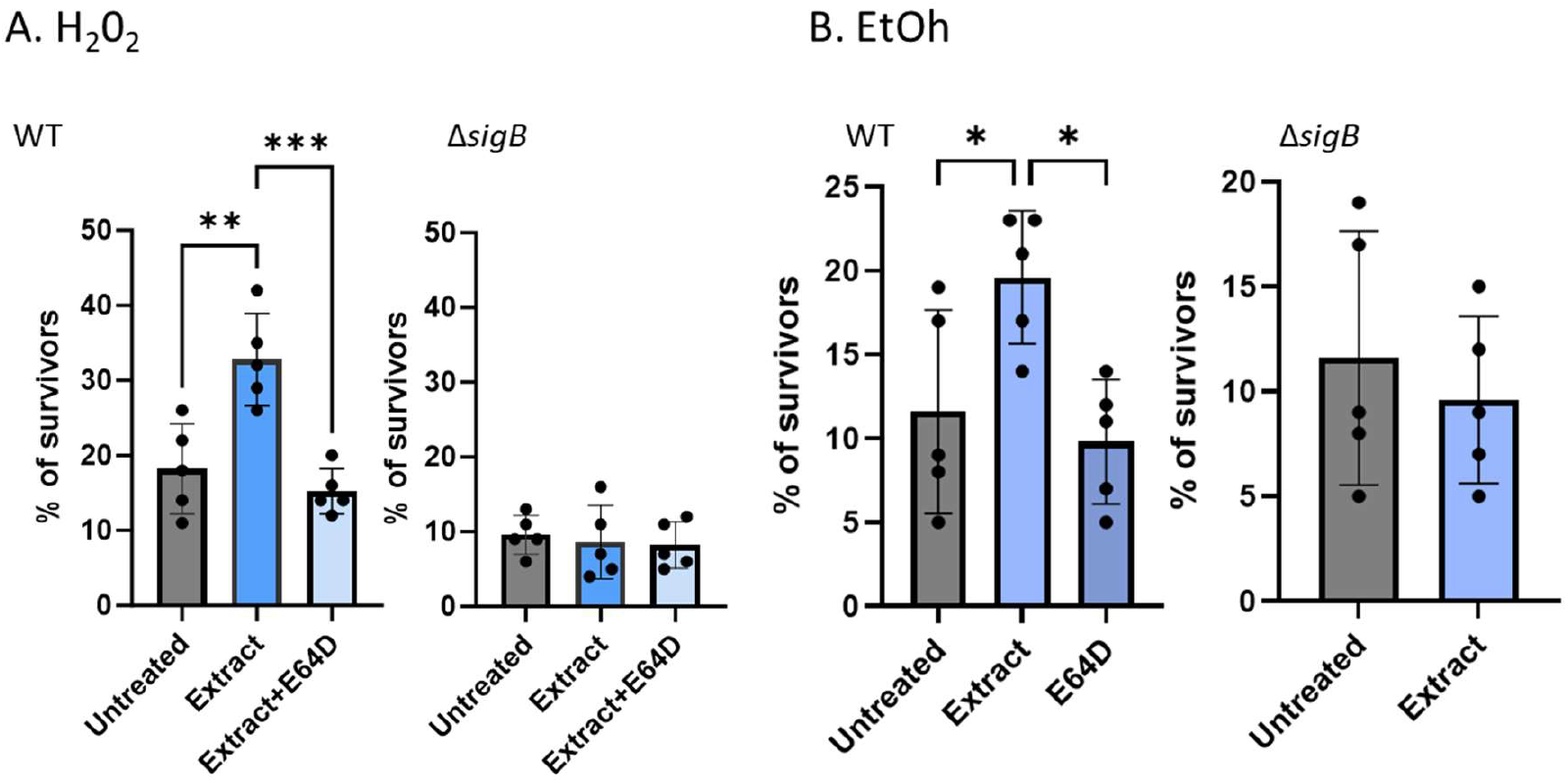
Amoeba CPs activity triggers resistance versus mild stressors. B. *subtilis NCIB3610* and its *ΔsigB* mutant were grown on MSgg agar at 30°C for 48 hours. Then, the colonies were cut in half with a razor blade. The 48-hour biofilms were re-suspended in 100μl PBS/Extract/Extract+E64D for 4 hours. Extracts were diluted v/v from a stock of 2ng/ μL. To determine the susceptibility to hydrogen peroxide or ethanol within the biofilms, cell-number percentage of CFU without or with chemical stress was compared. Cells from each group were pelleted and resuspended either in PBS 500μl or with hydrogen peroxide (10mM) or with 50% (v/v) ethanol (Gadot) for 10 minutes as indicated in each panel. Following incubation with stressors, biofilms were centrifuged (5 min at 14 000 r.p.m.), the supernatant was removed, and biofilms were resuspended in 500μl PBS and mildly sonicated (amplitude 20%, pulse 3 × 5 s). The number of CFU was determined by plating serial dilutions on LB plates and counting colonies after incubation at 30°C overnight. The percentage of surviving CFU is represented by the ratio of biofilm cells treated by the sterilizing agents compared with the same untreated group. Data represent the average and Standard deviation of five independent experiments performed in duplicates. *<0.05, **<0.01, and ***<0.001

### *B. subtilis* biofilm cells induce the cell wall stress response in response to predation

The protein TasA, targeted by CPs from *E. histolytica* for degradation, is in an intimate connection with the cell envelope, ^26^ and its degradation in pre-established biofilms may disturb the integrity of matrix-associated cell envelopes. Therefore, we tested whether cells from the partially degraded biofilms experience elevated stress following their exposure to the predator. Four sigma factors - SigM, SigW, SigX, and SigV - play crucial roles in maintaining cell envelope homeostasis, with SigW mutant having the most robust phenotype as a single mutant ^27^.

Our previous results demonstrated that SigW plays a central role in the activation of the β-lactamase PenP^28^ to provide resistance to nanomolar concentrations of β-lactam antibiotics ^29^. Exposing pre-established biofilms to amoeba extract significantly induced transcription from PenP promoter in a dose-dependent manner (Fig. 3A), although the level of induction was milder than the induction of the general stress response (Fig. 3B). The activation depended on the proteolytic activity of the amoeba’s CPs (see Fig. 3B). The transcription from *penP* promoter was not affected by the buffer or the presence of E64D (see Fig. S3). These results suggest that disrupting *B. subtilis* biofilms, rather than the presence of the amoeba’s CP and extract components, enhances the general stress response (see Fig. 1) and the cell wall stress response (see Fig. 3).

**Figure 3:**
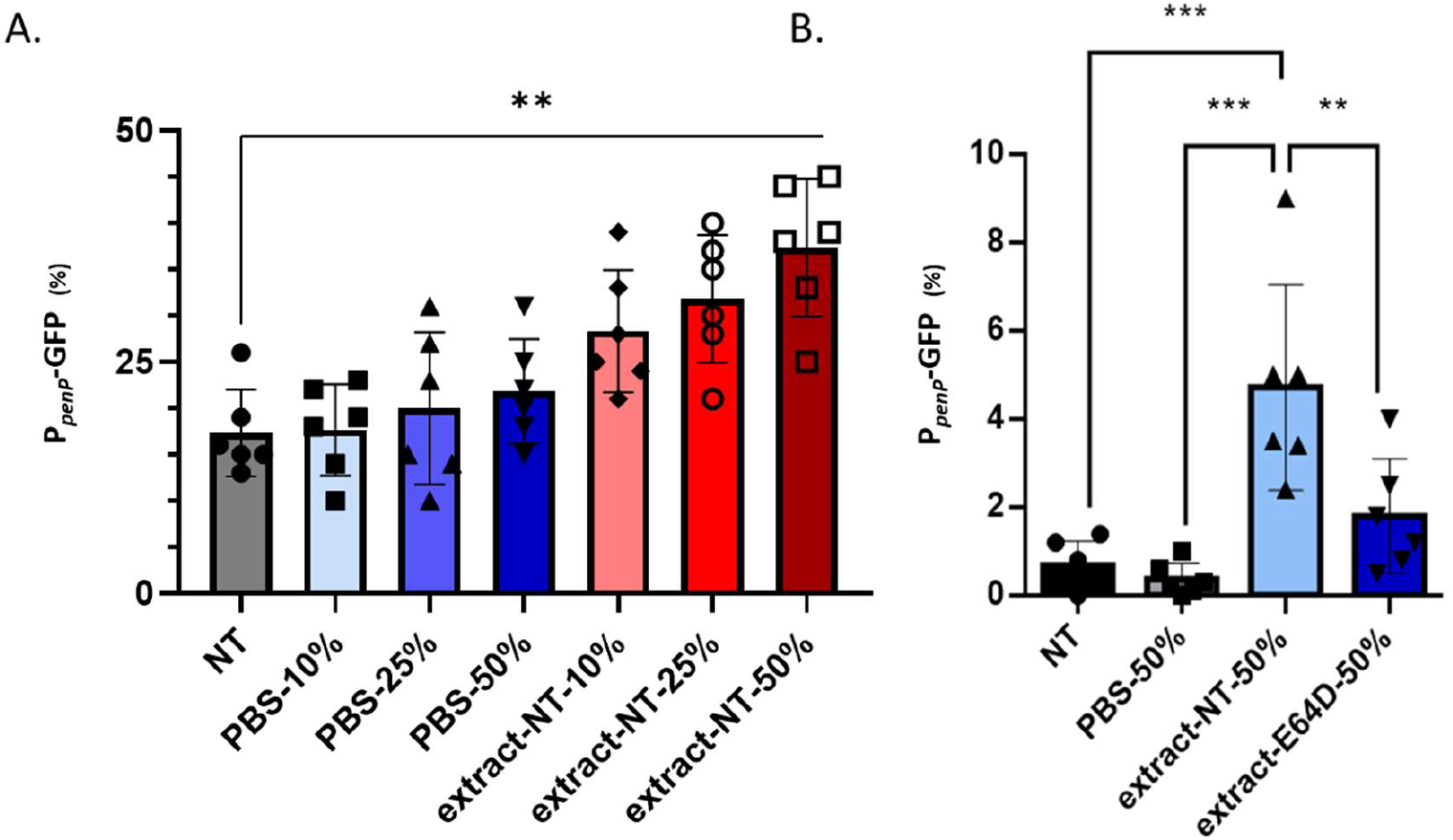
Amoeba CPs trigger the cell wall stress response. *B. subtilis* NCIB3610 wild type harboring *amyE::P*_*penP-*_*GFP* (Cell wall stress response) was analyzed in the presence and absence of increasing concentrations of E. histolytica’s extract using flow cytometry was analyzed in the presence and absence of increasing concentrations of *E. histolytica’s* extract using flow cytometry. Colonies were grown on MSgg medium and incubated at 30 °C. To determine the effect of amoeba’s lysate, the 48-hour biofilms were harvested. Colonies were incubated in indicated concentrations of PBS/Indicated concentrations of extract/ Indicated concentrations of extract+E64D for 4 hours. Extracts were diluted v/v from a stock of 2ng/ μL. From untreated and treated biofilms, 100,000 cells were counted. The % of cells expressing the reporters was calculated. Graphs represent mean ± SD from six independent experiments (n = 3). For B, extracts with and without E64D were used at identical concentrations (diluted v/v from a stock of 2ng/ μL). (B) As in A, except the colonies were re-suspended in 100μl PBS/Extract or PBS/Extract+E64D for 4 hours. Extracts were diluted v/v from a stock of 2ng/ μL. **<0.01, and ***<0.001.

Our previous results showed that due to the complete degradation of treated *B. subtilis* biofilms. The biofilm cells become sensitive to the treatment of toxic concentrations of Ampicillin (600 ng/ml Ampicillin) due to the breakdown of the extracellular matrix by CPs^17^. However, PenP only provided resistance to Ampicillin at low nanomolar concentrations. ^29^.

Therefore, we researched if activating PenP could safeguard cells impacted by the parasite when exposed to lower concentrations of Ampicillin (60 ng/ml Ampicillin). Our findings indicated a mild PenP-mediated protection against Ampicillin (Fig 4A and B), which relied on the activation of SigW (Fig. 4B). The effect of the *E. histolytica* extract appeared to be partly reliant on CPs (see Fig. 3B). In general, exposure to extract from *E. histolytica* had a more significant impact on the overall stress response and subsequent tolerance to related antibacterial stressors (e.g., H_2_O_2_) compared to the response to cell wall stress

**Figure 4:**
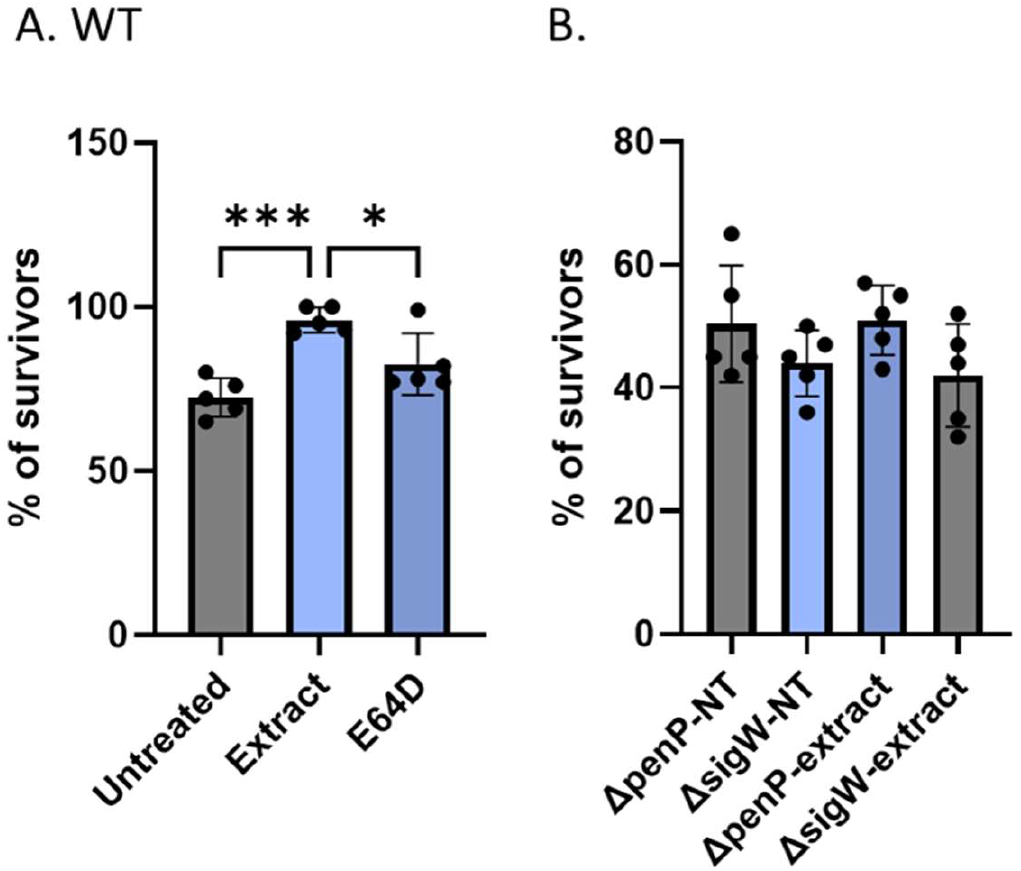
Amoeba CPs trigger the cell wall stress response. B. *subtilis NCIB3610* wild-type (A) and its indicated mutants (B) were grown on MSgg agar at 30°C for 48 hours. Then, the colonies were scrapped from the agar and re-suspended in 100μl PBS/Extract/Extract+E64D for 4 hours. Extracts were diluted v/v from an identical stock of 2ng/ μL in lysis buffer. Cells were pelleted and resuspended in PBS 500 μl with Ampicillin (60ng/ mL) for 4 hours. Following incubation, biofilm cells were centrifuged (5 min at 14 000 r.p.m.), the supernatant was removed, and biofilms were resuspended in 500μl PBS and mildly sonicated (amplitude 20%, pulse 3 × 5 s). The number of CFU was determined by plating serial dilutions on LB plates and counting colonies after incubation at 30°C overnight. The percentage of surviving CFU is represented by the ratio of biofilm cells treated by ampicillin, to their untreated counterpart. Data represent the average and Standard deviation of five independent experiments performed in duplicates. *<0.05, and ***<0.001.

### The exposure of planktonic cells to the extract reduces significantly the formation of biofilms

In planktonic cultures, the extract was toxic, in contrast to its lack of effect on biofilm cells. This toxicity was not dependent on the activity of CPs (Fig. S4). Consequently, we only assessed the impact of amoeba extract on expression in non-toxic conditions. Under these conditions, we did not observe any evidence of the activation of the SigB-dependent promoters of *ctc* and *katE* by the lysate of *E. histolytica*. These results suggest that pre-established biofilm cells respond to amoeba differently than planktonic bacteria (Fig. 5A and B). The response of biofilm cells was driven by the enzymatic activity of CPs, rather than by their presence (Figures 1 and 3). This effect may be less pronounced in the absence of an assembled extracellular protein matrix. Since it is not advantageous to form a biofilm in the presence of an efficient predator, we investigated whether a lysate from an amoeba predator could alter the dynamics of biofilm formation. To address this, we examined how the amoeba extract influences the expression of biofilm genes. Our findings revealed that the transcription from the tapA promoter, which controls the expression of the *tapA-sipW-tasA* operon ^30,31^ is significantly reduced in the presence of nanograms of amoeba extract (Fig. 5C). This result is consistent with the repression of biofilm formation with amoeba extract (Fig. 5D). Moreover, a similar repression was observed in the presence of E64D, but not with the buffer (Fig. 5C and S5). Interestingly, the repression of *tapA* promoter was not significant in treated biofilm cells (Fig. S6), further emphasizing a differential response of the biofilm cells and their planktonic counterparts.

**Figure 5:**
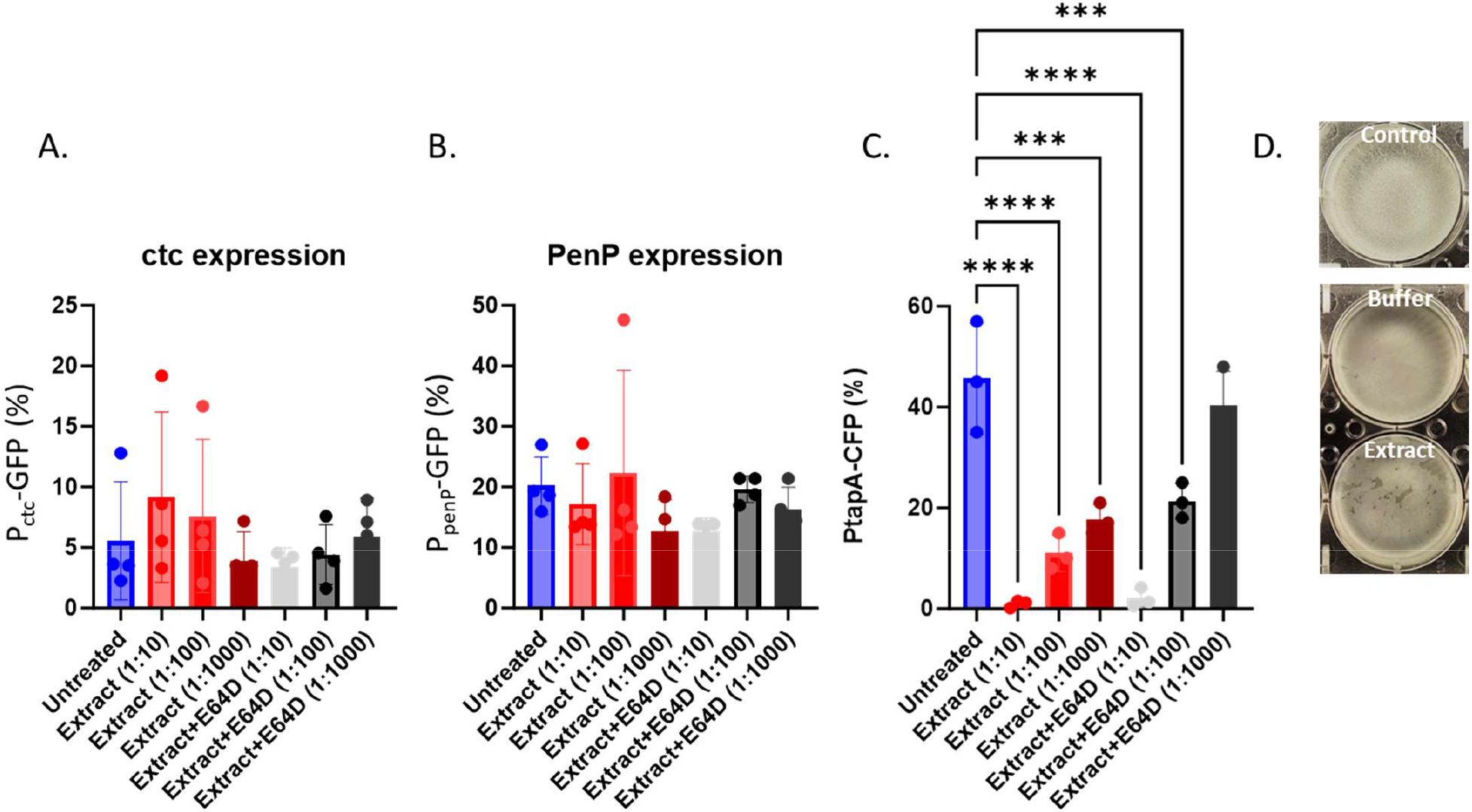
The response of planktonic cells to amoeba. (A-C) *B. subtilis* cells carrying each of the indicative reporters were grown logarithmically with shaking. At OD=0.6, cells were pelleted and re-suspended in 100μl in MSgg containing the indicated concentrations of PBS/Indicated concentrations of extract/ Indicated concentrations of extract+E64D for 4 hours. 100,000 cells were counted with flow cytometry from untreated and treated planktonic cells. The % of cells expressing the reporters was calculated. Graphs represent mean ± SD from four independent experiments performed with technical duplicates.). **<0.01, ***<0.001 and ****<0.0001. (D) Pellicle formation was assessed following 18 hours of growth in MSgg medium at 30 Celsius degrees either untreated (control) or with 0.1% buffer/extract as indicated.

## Discussion

*B. subtilis* biofilms engage in predator-prey interactions with protist parasites that can degrade pre-established biofilms pre-established biofilms ^17,18^. This degradation results from the specific activation of the amoeba’s cysteine proteinases (CPs) in the biofilm, which target and break down the biofilm’s protein matrix ^17^. The response of biofilm cells to changes in the microenvironment caused by parasites remains unclear. In this study, we demonstrate that the degradation of biofilms by amoeba’s CPs triggers a general stress response. This response involves the activation of a broad array of stress proteins, which are induced by various forms of physical stress, including heat, salt, ethanol, and acid stress. As a result, this stress response may serve a relatively nonspecific yet crucial protective function in the face of stress, regardless of the specific stressor. In our study, we have shown that the genes targeted by SigB are activated by the activity of amoeba’s CPs. Our conclusion is supported by the fact that the stress response induced by CPs is reversed in the presence of E64D. When the chemical inhibitor E64D is used, the CPs are present but inactive. This suggests that the dissolution of the biofilm imposes significant stress on the envelope of *B. subtilis*, leading to the activation of the extra cytoplasmic sigma factor SigW (as shown in Fig. 3 and 4). It’s worth noting that the stress signal is a result of the degradation of the biofilm rather than the presence of the CPs. This finding suggests that bacteria have evolved to perceive biofilm degradation as a stressor rather than simply a specific dispersal cue. Moreover, the differential response of bacterial cells at planktonic stage and biofilm stage is indicative of a fundamentally different defense strategy of the bacteria. As biofilm dissolves, the bacteria lose their physical and chemical protection by the extracellular matrix and require adaptation to potentially unsheltered environments^32,33^. The coupling of the activation of genes involved in stress protection with a general cue for biofilm degradation is beneficial to allow the bacteria surviving predation to increase their fitness while migrating to a less endangered environment (Fig. 6A).

**Figure 6:**
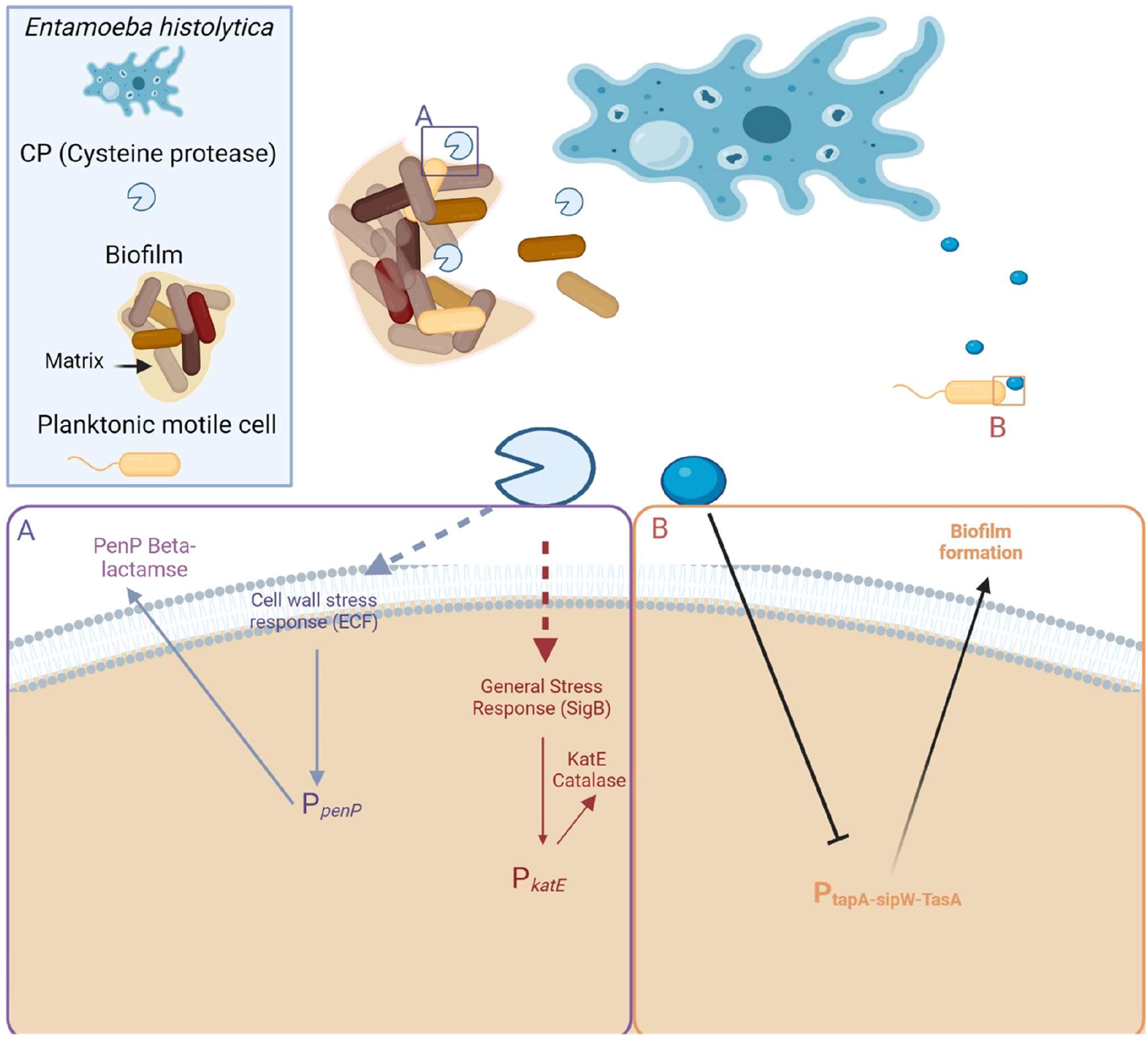
The response of biofilm and planktonic cells of B. subtilis to *Entamoeba histolytica*. The amoeba activates the general stress response and cell envelope cell response during biofilm degradation by CPs (A). In addition, the lysate also contains a signal repressing biofilm formation altogether in planktonic cells (B)

Interestingly, the response of planktonic bacteria to *E. histolytica* lysate suggests that planktonic bacteria sense and avoid biofilm formation in the presence of amoeba predators. The signal may be of relevance to trophozoites lysis during their rapid replication or while exposure to anti-parasitic drugs, or also be represented in their secretion or even volatiles’ profile (Fig. 6B).

Our discovery that *B. subtilis* biofilm genes are repressed in the presence of the parasite suggests a long-term arms race between the two. As bacteria detect the approaching predator, they inhibit the activation of biofilm genes-responsible for to the formation of a non-mobile community^12^ near the parasite. Downregulation of biofilm formation allows the bacteria to migrate toward alternative environments. Interestingly, the differential responses of planktonic bacterial cells and biofilm cells indicate that the differentiation into a multicellular community also influences their responses to predators and aggressors.

Collectively, our results suggest that the antagonistic coevolution between biofilm prey and protist parasites may significantly impact the evolution of bacterial transcriptional regulation.

## References

1. Haque, R., Huston, C.D., Hughes, M., Houpt, E., and Petri, W.A., Jr. (2003). Amebiasis. N Engl J Med 348, 1565–1573. 10.1056/NEJMra022710348/16/1565 [pii].

2. Guillen, N. (2021). Signals and signal transduction pathways in Entamoeba histolytica during the life cycle and when interacting with bacteria or human cells. Molecular microbiology 115, 901–915. 10.1111/mmi.14657.

3. Vlamakis, H., Chai, Y., Beauregard, P., Losick, R., and Kolter, R. (2013). Sticking together: building a biofilm the Bacillus subtilis way. Nature reviews. Microbiology 11, 157–168. 10.1038/nrmicro2960.

4. Karatan, E., and Watnick, P. (2009). Signals, regulatory networks, and materials that build and break bacterial biofilms. Microbiol Mol Biol Rev 73, 310–347. 10.1128/MMBR.00041-08.

5. Romero, D., Aguilar, C., Losick, R., and Kolter, R. (2010). Amyloid fibers provide structural integrity to Bacillus subtilis biofilms. Proc Natl Acad Sci U S A 107, 2230–2234. 10.1073/pnas.0910560107.

6. Azulay, D.N., Spaeker, O., Ghrayeb, M., Wilsch-Brauninger, M., Scoppola, E., Burghammer, M., Zizak, I., Bertinetti, L., Politi, Y., and Chai, L. (2022). Multiscale X-ray study of Bacillus subtilis biofilms reveals interlinked structural hierarchy and elemental heterogeneity. Proc Natl Acad Sci U S A 119. 10.1073/pnas.2118107119.

7. Romero, D., Vlamakis, H., Losick, R., and Kolter, R. (2014). Functional analysis of the accessory protein TapA in Bacillus subtilis amyloid fiber assembly. J Bacteriol 196, 1505–1513. 10.1128/JB.01363-13.

8. Camara-Almiron, J., Dominguez-Garcia, L., El Mammeri, N., Lends, A., Habenstein, B., de Vicente, A., Loquet, A., and Romero, D. (2023). Molecular characterization of the N-terminal half of TasA during amyloid-like assembly and its contribution to Bacillus subtilis biofilm formation. NPJ biofilms and microbiomes 9, 68. 10.1038/s41522-023-00437-w.

9. Camara-Almiron, J., Navarro, Y., Diaz-Martinez, L., Magno-Perez-Bryan, M.C., Molina-Santiago, C., Pearson, J.R., de Vicente, A., Perez-Garcia, A., and Romero, D. (2020). Dual functionality of the amyloid protein TasA in Bacillus physiology and fitness on the phylloplane. Nature communications 11, 1859. 10.1038/s41467-020-15758-z.

10. Kobayashi, K., and Iwano, M. (2012). BslA(YuaB) forms a hydrophobic layer on the surface of Bacillus subtilis biofilms. Molecular microbiology 85, 51–66. 10.1111/j.1365-2958.2012.08094.x.

11. Arnaouteli, S., Bamford, N.C., Stanley-Wall, N.R., and Kovacs, A.T. (2021). Bacillus subtilis biofilm formation and social interactions. Nature reviews. Microbiology 19, 600–614. 10.1038/s41579-021-00540-9.

12. Steinberg, N., Keren-Paz, A., Hou, Q., Doron, S., Yanuka-Golub, K., Olender, T., Hadar, R., Rosenberg, G., Jain, R., Camara-Almiron, J., et al. (2020). The extracellular matrix protein TasA is a developmental cue that maintains a motile subpopulation within Bacillus subtilis biofilms. Science signaling 13. 10.1126/scisignal.aaw8905.

13. Vlamakis, H., Aguilar, C., Losick, R., and Kolter, R. (2008). Control of cell fate by the formation of an architecturally complex bacterial community. Genes & development 22, 945–953. 10.1101/gad.1645008.

14. Aguilar, C., Vlamakis, H., Guzman, A., Losick, R., and Kolter, R. (2010). KinD is a checkpoint protein linking spore formation to extracellular-matrix production in Bacillus subtilis biofilms. mBio 1. 10.1128/mBio.00035-10.

15. Keren-Paz, A., Brumfeld, V., Oppenheimer-Shaanan, Y., and Kolodkin-Gal, I. (2018). Micro-CT X-ray imaging exposes structured diffusion barriers within biofilms. NPJ biofilms and microbiomes 4, 8. 10.1038/s41522-018-0051-8.

16. Pane-Farre, J., Quin, M.B., Lewis, R.J., and Marles-Wright, J. (2017). Structure and Function of the Stressosome Signalling Hub. Subcell Biochem 83, 1–41. 10.1007/978-3-319-46503-6_1.

17. Zanditenas, E., Trebicz-Geffen, M., Kolli, D., Dominguez-Garcia, L., Farhi, E., Linde, L., Romero, D., Chapman, M., Kolodkin-Gal, I., and Ankri, S. (2023). Digestive exophagy of biofilms by intestinal amoeba and its impact on stress tolerance and cytotoxicity. NPJ biofilms and microbiomes 9, 77. 10.1038/s41522-023-00444-x.

18. Zanditenas, E., and Ankri, S. (2024). Unraveling the interplay between unicellular parasites and bacterial biofilms: Implications for disease persistence and antibiotic resistance. Virulence 15, 2289775. 10.1080/21505594.2023.2289775.

19. Rigothier, M.C., Khun, H., Tavares, P., Cardona, A., Huerre, M., and Guillen, N. (2002). Fate of Entamoeba histolytica during establishment of amoebic liver abscess analyzed by quantitative radioimaging and histology. Infect Immun 70, 3208–3215. 10.1128/IAI.70.6.3208-3215.2002.

20. Dupuy, M., Berne, F., Herbelin, P., Binet, M., Berthelot, N., Rodier, M.H., Soreau, S., and Hechard, Y. (2014). Sensitivity of free-living amoeba trophozoites and cysts to water disinfectants. Int J Hyg Environ Health 217, 335–339. 10.1016/j.ijheh.2013.07.007.

21. Diamond, L.S., Harlow, D.R., and Cunnick, C.C. (1978). A new medium for the axenic cultivation of Entamoeba histolytica and other Entamoeba. Trans R Soc Trop Med Hyg 72, 431–432. 10.1016/0035-9203(78)90144-x.

22. Branda, S.S., Gonzalez-Pastor, J.E., Ben-Yehuda, S., Losick, R., and Kolter, R. (2001). Fruiting body formation by Bacillus subtilis. Proc Natl Acad Sci U S A 98, 11621–11626. 10.1073/pnas.191384198.

23. Maan, H., Itkin, M., Malitsky, S., Friedman, J., and Kolodkin-Gal, I. (2022). Resolving the conflict between antibiotic production and rapid growth by recognition of peptidoglycan of susceptible competitors. Nature communications 13, 431. 10.1038/s41467-021-27904-2.

24. Waters, S.M., Robles-Martinez, J.A., and Nicholson, W.L. (2014). Exposure of Bacillus subtilis to low pressure (5 kilopascals) induces several global regulons, including those involved in the SigB-mediated general stress response. Applied and environmental microbiology 80, 4788–4794. 10.1128/AEM.00885-14.

25. Petersohn, A., Engelmann, S., Setlow, P., and Hecker, M. (1999). The katX gene of Bacillus subtilis is under dual control of sigmaB and sigmaF. Mol Gen Genet 262, 173–179. 10.1007/s004380051072.

26. Romero, D., Vlamakis, H., Losick, R., and Kolter, R. (2011). An accessory protein required for anchoring and assembly of amyloid fibres in B. subtilis biofilms. Molecular microbiology 80, 1155–1168. 10.1111/j.1365-2958.2011.07653.x.

27. Helmann, J.D. (2016). Bacillus subtilis extracytoplasmic function (ECF) sigma factors and defense of the cell envelope. Curr Opin Microbiol 30, 122–132. 10.1016/j.mib.2016.02.002.

28. Imanaka, T., Oshihara, W., Himeno, T., and Aiba, S. (1983). Comparative studies on extracellular penicillinases of the same structural gene, penP, expressed in Bacillus licheniformis and Bacillus subtilis. J Gen Microbiol 129, 2621–2628. 10.1099/00221287-129-8-2621.

29. Bucher, T., Keren-Paz, A., Hausser, J., Olender, T., Cytryn, E., and Kolodkin-Gal, I. (2019). An active beta-lactamase is a part of an orchestrated cell wall stress resistance network of Bacillus subtilis and related rhizosphere species. Environmental microbiology 21, 1068–1085. 10.1111/1462-2920.14526.

30. Kearns, D.B., Chu, F., Branda, S.S., Kolter, R., and Losick, R. (2005). A master regulator for biofilm formation by Bacillus subtilis. Molecular microbiology 55, 739–749. 10.1111/j.1365-2958.2004.04440.x.

31. Chu, F., Kearns, D.B., Branda, S.S., Kolter, R., and Losick, R. (2006). Targets of the master regulator of biofilm formation in Bacillus subtilis. Molecular microbiology 59, 1216–1228. 10.1111/j.1365-2958.2005.05019.x.

32. Steinberg, N., and Kolodkin-Gal, I. (2015). The Matrix Reloaded: Probing the Extracellular Matrix Synchronizes Bacterial Communities. J Bacteriol 197, 2092–2103. 10.1128/JB.02516-14.

33. Karygianni, L., Ren, Z., Koo, H., and Thurnheer, T. (2020). Biofilm Matrixome: Extracellular Components in Structured Microbial Communities. Trends in microbiology 28, 668–681. 10.1016/j.tim.2020.03.016.

