## Supporting Information for "Differential coping strategies exerted by biofilm and planktonic cells of the beneficial bacterium *B. subtilis* in response to the protozoan predator *Entamoeba histolytica*"

### Supporting figures

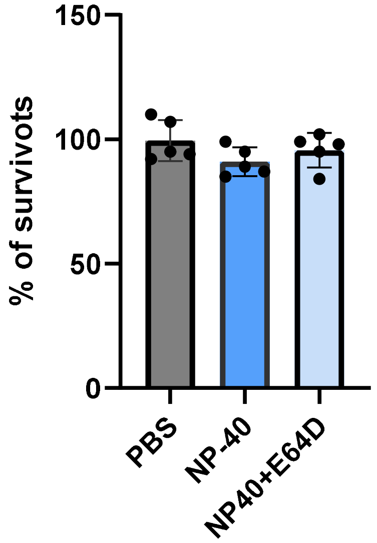


Figure S1: **Monitoring the effect of lysis buffer on biofilm cells.**

*B. subtilis* NCIB3610 harboring *amyE::P*$_{ctc}$*-GFP* (General stress response) was analyzed in the presence and absence of increasing concentrations of NP-40 lysis buffer and E64D. The 48-hour biofilms were re- suspended in 100µl of indicated solution representing 50% extract working concentrations: (0.2% NP-40, with/without E64D 5 µM)

Following incubation with stressors, biofilms were centrifuged (5 min at 14 000 r.p.m.), the supernatant was removed, and biofilms were resuspended in 500µl PBS and mildly sonicated (amplitude 20%, pulse 3 × 5 s). The number of CFU was determined by plating serial dilutions on LB plates and counting colonies after incubation at 30°C overnight. The percentage of surviving CFU is represented by the ratio of biofilm cells treated by the sterilizing agents compared with the same untreated group.  Data represent the average and Standard deviation of five independent experiments performed in duplicates.

No significant difference was observed following statistical analysis.

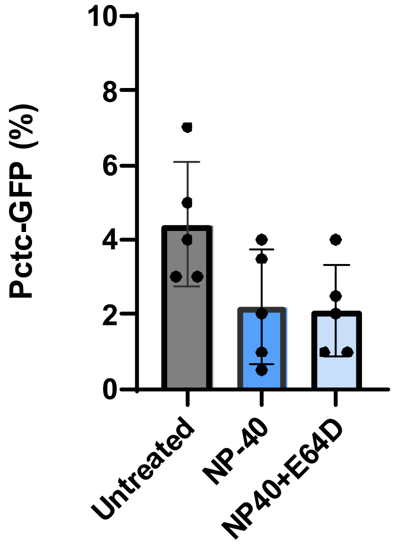


Figure S2: **Monitoring the effect of lysis buffer on transcription from the *ctc* promoter.**

*B. subtilis* NCIB3610 harboring *amyE::P$_{ctc}$-GFP* (General stress response) was analyzed in the presence and absence of increasing concentrations of NP-40 lysis buffer and E64D. The 48-hour biofilms were re- suspended in 100µl of indicated solution representing 50% extract working concentrations: (0.2% NP-40, with/without E64D 5 µM). Flow cytometry was performed as described in Figure 2. No significant difference was observed following statistical analysis.

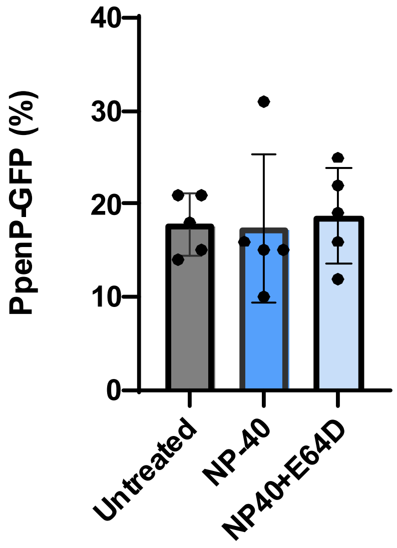


Figure S3: **Monitoring the effect of lysis buffer on transcription from the *penP* promoter.**

*B. subtilis* NCIB3610 harboring *amyE::P$_{penP}$-GFP* was analyzed in the presence and absence of increasing concentrations of NP-40 lysis buffer an:d E64D. The 48-hour biofilms were re-suspended in 100µl of indicated solution representing 50% extract working concentrations: (0.2% NP-40, with/without E64D 5 µM). Flow cytometry was performed as described in Figure 2. No significant difference was observed following statistical analysis.

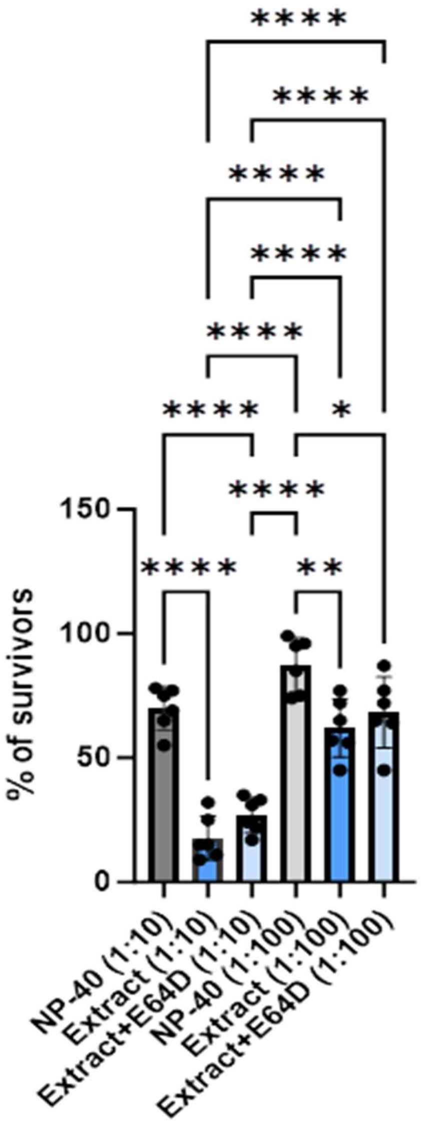


Figure S4: **The response of planktonic cells to *E. histolytica* extract.** B. subtilis cells carrying the indicative reporters were grown logarithmically with shaking. At OD=0.6, cells were pelleted and re- suspended in 100µl in MSgg containing the indicated concentrations of PBS+NP40/Indicated concentrations of extract/ Indicated concentrations of extract+E64D for 4 hours from a stock of 2ng/ µL. Following incubation with stressors, cells were centrifuged (5 min at 14 000 r.p.m.), the supernatant was removed, and biofilms were resuspended in 500µl PBS.

The number of CFU was determined by plating serial dilutions on LB plates and counting colonies after incubation at 30°C overnight. The percentage of surviving CFU is represented by the ratio of biofilm cells treated by the sterilizing agents compared with the same untreated group. Data represent the average and Standard deviation of five independent experiments performed in duplicates. *<0.05, **<0.01, ***<0.001 and ****<0.0001

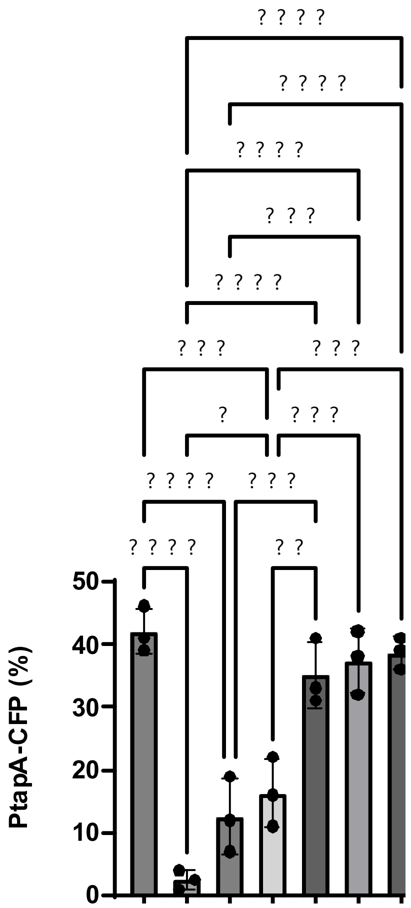


Figure S5: **The transcription from the *tapA* promoter in the presence of the *E. histolytica* extract.**

*B. subtilis* cells carrying the *tapA* transcriptional reporter were grown logarithmically with shaking. At OD=0.6, cells were pelleted and re- suspended in 100µl in MSgg containing the indicated concentrations of PBS+NP40/Indicated concentrations of extract from a stock of 2ng/ µL. Following incubation with stressors, cells were centrifuged (5 min at 14 000 r.p.m.), the supernatant was removed, and biofilms were resuspended in 500µl PBS. From untreated and treated biofilms, 100,000 cells were counted with flow cytometry. The % of cells expressing the reporters was calculated. Graphs represent mean ± SD from six independent experiments (n = 2). **<0.01, ***<0.001 and ****<0.0001.

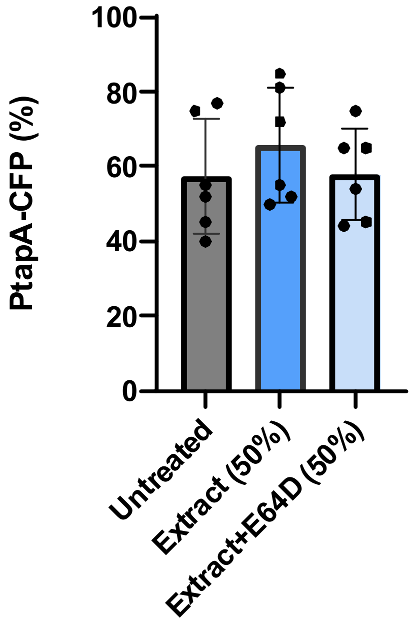


Figure S6: **The transcription from the *tapA* promoter is not sensitive to E64D**

*B. subtilis* cells carrying the *tapA* transcriptional reporter were grown logarithmically with shaking. At OD=0.6, cells were pelleted and re- suspended in 100µl in MSgg (untreated)/ MSgg with extract/MSGG with extract and E64D from a stock of 2ng/ µL. Following incubation with stressors, cells were centrifuged (5 min at 14 000 r.p.m.), the supernatant was removed, and biofilms were resuspended in 500µl PBS. From untreated and treated biofilms, 100,000 cells were counted with flow cytometry. The % of cells expressing the reporters was calculated. Graphs represent mean ± SD from six independent experiments (n = 2). No significant difference was observed following statistical analysis.
